# Endoparasites and vector-borne pathogens in dogs from Greek islands: pathogen distribution and zoonotic implications

**DOI:** 10.1101/472365

**Authors:** Anastasia Diakou, Angela Di Cesare, Simone Morelli, Mariasole Colombo, Lenaig Halos, Giulia Simonato, Androniki Tamvakis, Frederic Beugnet, Barbara Paoletti, Donato Traversa

## Abstract

The present study investigated the presence of endo- and ecto-parasites, and vector-borne pathogens, in dogs from four islands of Greece. A total of 200 owned and sheltered dogs were examined with different microscopic, serological and molecular methods.

Of the examined dogs, 130 (65%) were positive for one or more parasites and/or vector-borne pathogens. The most common zoonotic intestinal helminths recorded were Ancylostomatidae (12.5%) and *Toxocara canis* (3.5%). Ninety-three dogs (46.5%) seroreacted to *Rickettsia conorii*. Twenty-two (11%) of them were also PCR positive and 7 (3.5%) showed corpuscoles suggestive of *Rickettsia* spp. on the blood smears. Nineteen dogs (9.5%) were seropositive for *Ehrlichia canis*, three of them being also PCR positive. Dogs positive for *Anaplasma phagocytophilum-Anaplasma platys* (1%), *Dirofilaria immitis* (0.5%) and *Babesia canis* (0.5%) were also found. Fleas and ticks were recorded in 53 (26.5%) and 50 (25%) dogs and all specimens were identified as *Ctenocephalides felis felis* and *Rhipicephalus sanguineus sensu latu*. Binary multiple univariate Generalized Linear Models were used to investigate factors and clinical signs related to the recorded positivity, while the association of specific signs with the pathogens was evaluated using tests of independence. Knowledge of occurrence and impact of zoonotic parasites and vector-borne pathogens in dog populations is crucial to prevent the infection in animals and people, and to control the risk of spreading of these pathogens in endemic and non-endemic areas.

**Author summary:** Both owned and sheltered dogs can harbor a variety of intestinal and extra-intestinal endoparasites, as well as vector-borne pathogens and ectoparasites, of zoonotic concern. Dog shelters and stray dogs are present in several touristic areas of Greece, including Sporades and Cyclades islands, where tourists often bring their pets with them, likely travelling from non-endemic to endemic areas. The present study has been carried out with the aim to evaluate the occurrence of the aforementioned pathogens. Data obtained showed that they are present in canine populations of Greece, with possibilities of infection for travelling dogs, which can also contribute to the spreading of zoonotic vector-borne diseases, introducing new pathogens in previously non-endemic areas. For these reasons, a constant monitoring of the epidemiological situation, improving control measures and correct diagnostic approaches are of primary importance for the prevention of canine and human infections, decreasing the spreading of potentially deadly pathogens.

## Introduction

Several parasitoses (e.g. internal helminthoses) and vector-borne diseases (VBDs) of veterinary importance represent a serious hazard for human health, particularly when transmission pressure and circulation of zoonotic infections are difficult to control. Because of a lifestyle that implies a low-grade of sanitary management, stray and free-roaming dogs are at high risk of becoming infected with a wide range of pathogens. Consequently, they act as a permanent source of infection for vectors, other animals and humans [1, 2].

Although dogs are efficient sentinels for investigating the occurrence and the epidemiological impact of zoonoses [2, 3] and a constant sanitary monitoring of canine populations is crucial, data on the simultaneous occurrence of endo/ecto-parasites and VBDs in several Mediterranean areas are still limited to specific narrow areas or to selected pathogens [4–7].

The number of stray dogs in most Greek Islands is low but many privately owned animals have a free-ranging lifestyle due to the local rural territory. Moreover, some islands have shelters for stray and abandoned animals, where prevention and treatment regimens are not regularly applied. At the same time, the number of dogs travelling to and from the insular Greece is increasing in proportion to tourism [8] with the realistic risk that these pets may acquire or introduce pathogens. Interestingly, a recent study conducted in Greece has shown that cats may be infected and/or exposed to a number of intestinal parasites and agents transmitted by arthropods, several of them of zoonotic importance and potentially shared between cats and dogs [9]. Therefore, the present study aimed to investigate the simultaneous occurrence of intestinal parasites, vector-borne pathogens and ectoparasites, in different canine populations, including stray and owned dogs in four regions of insular Greece.

## Methods

### Study areas and sampling

The study was conducted in four islands of Greece. Authorizations to sample and examine the dogs were obtained case-by-case from local authorities, animal responsibles/owners and animal rights organisations. Available data on sex, breed, living conditions, age and presence or absence of clinical signs compatible with parasitoses were registered for each animal. Both faecal and blood samples were collected from dogs, that were also examined for fleas, ticks, ear mites and, in the presence of compatible skin lesions, body manges.

Ethics approval is not applicable as all activities carried out on dogs in the present study have been perfrormed by routine diagnostics procedures performed by veterinarians working on each study site. Consent to examine and sample the animals have been obtained, case by case, from local authorities, animal owners and animal right organisations. Therefore, no official ethics permission or further authorization was required.

### Copromicroscopic examinations

Faecal samples were examined using standard zinc sulphate flotation and merthiolate iodine formaldehyde (MIF) - ether sedimentation methods [10, 11].

### Blood analysis

#### Microscopic examinations

Giemsa stained blood smears were performed to evaluate by light microscopy at 1000× magnification the presence of *Babesia* spp., *Rickettsia* spp., *Ehrlichia* spp. and *Anaplasma* spp. elements, based on morphology, size and cellular tropism [12, 13].

A modified Knott’s technique was performed to detect and identify circulating microfilariae under light microscope (100×, 200× and 400× magnifications) [12]. If present, microfilariae were identified on the basis of differential morphometric (i.e. length and width) and morphological (i.e. anterior and posterior end) characteristics [14, 15].

#### Serological examinations

Sera obtained from blood samples were subjected to the following serological examinations, according to the manufacturers’ instructions.

-SNAP 4Dx Plus test (IDEXX Laboratories, Inc. USA) for the detection of *Dirofilaria immitis* circulating antigens and of antibodies against *Anaplasma phagocytophilum/Anaplasma platys, Ehrlichia canis* and *Borrelia burgdorferi*;

-*Leishmania* IC test (Agrolabo diagnostics, Italy) for the qualitative detection of anti-*L. infantum* antibodies;

-indirect immunofluorescence antibody assay kits (IFAT) “Mega FLUO^®^ BABESIA *canis*” (Megacor Diagnostik GmbH, Austria) and “Mega FLUO^®^ RICKETTSIA *conorii*” (Megacor Diagnostik GmbH, Austria) for the detection of *anti-Babesia canis-IgG* and *anti-Rickettsia conorii*-IgG, with a screening dilution of 1:128 and 1:64 for *B. canis* and *R. conorii* respectively.

Positive and negative control sera were included in each test series.

#### Molecular detection

Genomic DNA was extracted from blood samples using a commercial kit (QIAamp DNA blood Mini kit - Qiagen GmbH, Hilden, Germany), and PCR positivity to Anaplasmatacea, *Rickettsia* spp., *Babesia* spp. and *Leishmania* spp. DNA was tested by different protocols (Table 1) [16, 17].

**Table 1.**
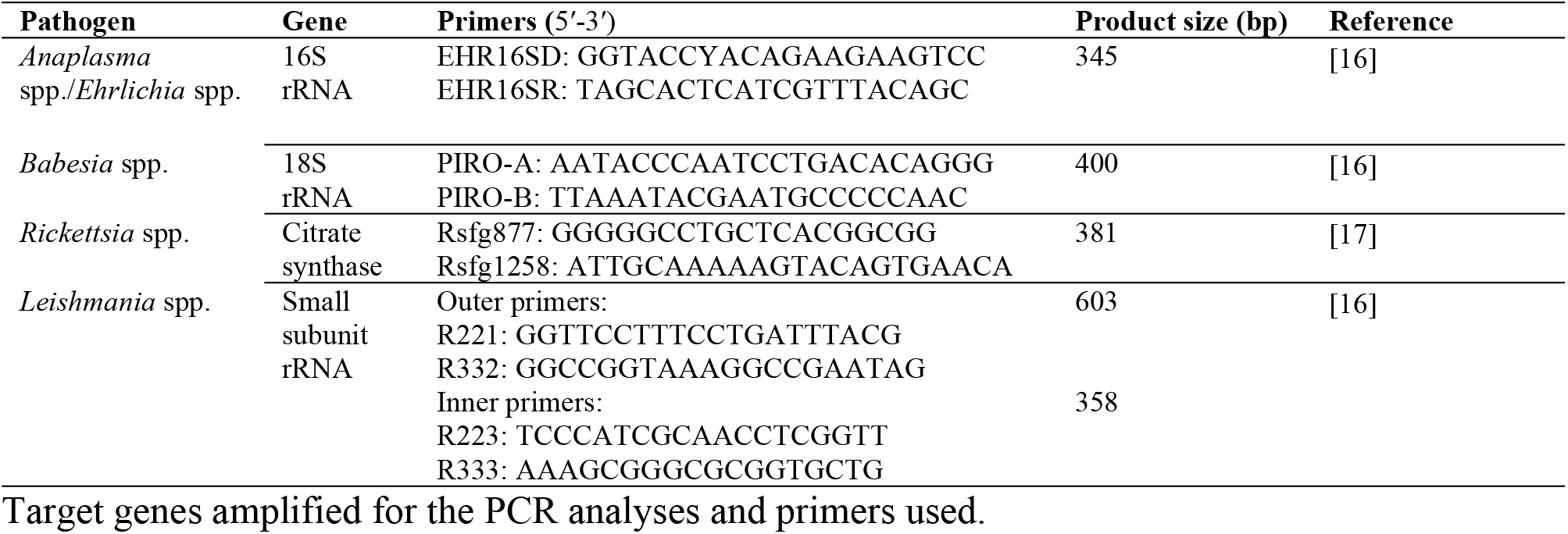

PCR products were individually sequenced directly in an automated sequencer. Sequences were determined in both directions, aligned using ClustalX software and analyzed with sequences available in GenBankTM using Nucleotide Basic Local Alignment Search Tool (BLASTN) [18].

### Ectoparasites

#### Fleas and ticks

The entire body of each dog was examined with an extra-fine flea comb. Fleas seen on the comb were removed using forceps and placed in a 2 ml microtube containing 70% ethanol for storage. If fleas were not present, any debris found on the comb was transferred to a piece of moist white paper, and animals were considered infected if the debris dissolved into a red color.

Ticks were individuated by thumb-counting, removed using forceps and placed in a 2 ml microtube containing 70% ethanol pending identification.

Collected fleas and ticks were identified by standard morphologic and morphometric keys [11, 19, 20].

#### Mites

All dogs were subjected to an otoscopic examination by ear swabs to detect the presence of any sign (e.g. errhytema, black/brown waxy discharge) caused by the ear mite *Otodectes cynotis*. Each ear swab was smeared with the addition of small quantity of mineral oil onto a glass slide, and examined under a microscope to identify mites by standard morphological keys [11, 21]. In the presence of compatible clinical signs, e.g. skin areas with loss of hair, itching, reddened rash, yellowish crusts, dogs were examined for manges as deemed appropriate. The collected material was examined under a microscope to identify mites by standard morphological keys [11, 21].

### Statistical analysis

Statistical analysis was performed to evaluate the association of five main factors (study site, age, sex, lifestyle and travel history) with infections and parasitoses detected in the examined dogs, especially with zoonotic potential. Binary multiple univariate Generalized Linear Models (GLM) were used to test the above mentioned factors with the presence of endoparasites (intestinal helminths and filariae), exposure/positivity for VBDs, positivity to zoonotic pathogens (*R. conorii, Anaplasma* spp., *L. infantum, Dirofilaria* spp., Ancylostomatidae and *T. canis*) and presence of ectoparasites [22]. Furthermore, the determined odds ratio (OR) was used to measure the strength of association between the values of each factor to the presence of each infection. The same analysis was applied to test whether the occurrence of four major clinical signs (i.e. dermatological signs, ocular manifestations, weight loss and pale mucosae) was related to the detection of a VBD.

Finally, the association of gastrointestinal signs with intestinal parasitism and ectoparasitoses as evidence of exposure to VBDs was statistically tested using Fisher’s exact test of independence [23]. The statistical analysis was implemented using the R package version 3.2.2 (R Development Core Team, 2006).

## Results

### Enrolment and geographical distribution

Overall 200 dogs, i.e. 66, 50, 43 and 41 dogs from four islands, Santorini, Tinos, Ios and Skiathos, were included in the survey, respectively.

Table 2 reports age, sex and lifestyle of study dogs. In particular, 87 were privately owned with no travel history, 36 were pets that had travelled with their owners at least once (i.e. 32 across Greece, 2 in Bulgaria, 1 in Italy and 1 in different European countries), and 77 where sheltered animals. Both owned and sheltered dogs were highly distributed within different islands and age groups, with the exception of stray animals less than 1-year-old that were found only in Santorini (n=13). Nevertheless, the dataset was considered as sufficiently large (n=200) so that the heuristic rule (i.e. a few of the expected cells counts less than five is not violated) provides safe statistical results [24].

**Table 2.**
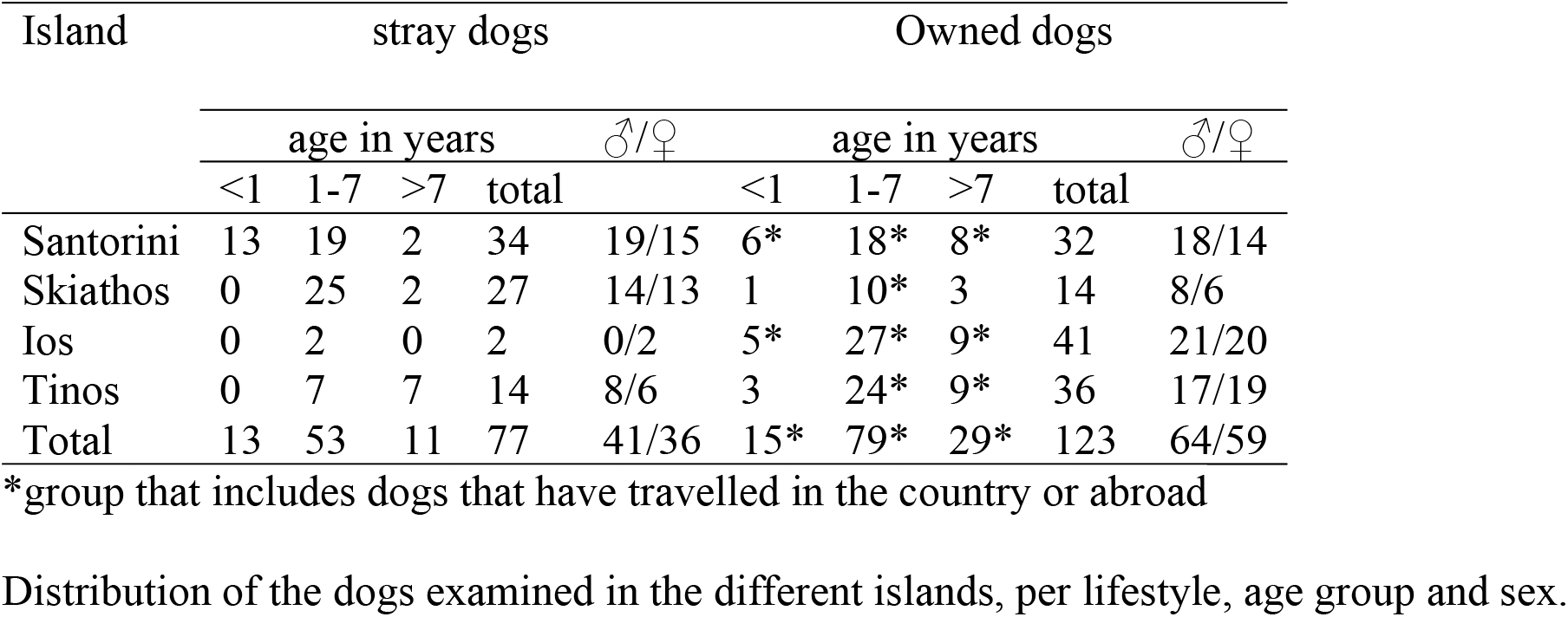

| Island | stray dogs |  |  |  |  | Owned dogs |  |  |  |  |
| --- | --- | --- | --- | --- | --- | --- | --- | --- | --- | --- |
|  | age in years |  |  |  | ♂/♀ | age in years |  |  |  | ♂/♀ |
|  | <1 | 1-7 | >7 | total |  | <1 | 1-7 | >7 | total |  |
| Santorini | 13 | 19 | 2 | 34 | 19/15 | 6* | 18* | 8* | 32 | 18/14 |
| Skiathos | 0 | 25 | 2 | 27 | 14/13 | 1 | 10* | 3 | 14 | 8/6 |
| Ios | 0 | 2 | 0 | 2 | 0/2 | 5* | 27* | 9* | 41 | 21/20 |
| Tinos | 0 | 7 | 7 | 14 | 8/6 | 3 | 24* | 9* | 36 | 17/19 |
| Total | 13 | 53 | 11 | 77 | 41/36 | 15* | 79* | 29* | 123 | 64/59 |

\*group that includes dogs that have travelled in the country or abroad
Distribution of the dogs examined in the different islands, per lifestyle, age group and sex.

### Exposure to parasites, vector borne diseases and prevalence of zoonotic agents

Overall 130 dogs (65%) were positive for at least one intestinal parasite or VBD. In particular, 96 (48%) showed a monospecific infection and 34 (17%) scored positive for mixed infections by endoparasites and VBD. Two animals (1%) were infected by more than 1 endoparasite, 16 (8%) by more than one VBD and 16 (8%) were infected by both intestinal parasites and VBDs. Various zoonotic agents were found in 128 (64%) dogs. Fleas and ticks were recorded in 53 (26.5%) and 50 (25%) dogs respectively.

#### Endoparasites

Overall thirty-six (18%) dogs were positive for at least one endoparasite at copromicroscopic examinations. Ancylostomatidae (12.5%, 25/200) and *Toxocara canis* (i.e. 3.5%, 7/200) were the most common zoonotic helminths. Non-zoonotic infections by *Trichuris vulpis* (3.5%) and coccidia (1%) were also found (Table 3). Eggs of the zoonotic rat tapeworm *Hymenolepis diminuta* were detected in the faeces of one dog, while eggs of the trematode *Dicrocoelium dendriticum* were found in the faeces of another animal.

The relative prevalence of intestinal parasites was related to the island of residence, the age and the lifestyle of the animal (Table 4). In particular, the odd of intestinal parasites occurrence was 5.49, 11.43 and 23.86 times higher in animals residing in Santorini, compared to those that reside in Tinos, Ios and Skiathos, respectively. Animals ageing 1 to 7 years were found to be 4 times (i.e. 1/0.25) more likely to be infected compared to young animals up to 1 year of age. Furthermore, a higher risk of intestinal parasites prevalence was found in stray dogs (odds ratio=11.37) compared to owned dogs. Nonetheless, the overall occurence of intestinal parasites was not related to the presence of clinical signs (Fisher’s exact test, *p*= 0.3283).

**Table 3.**
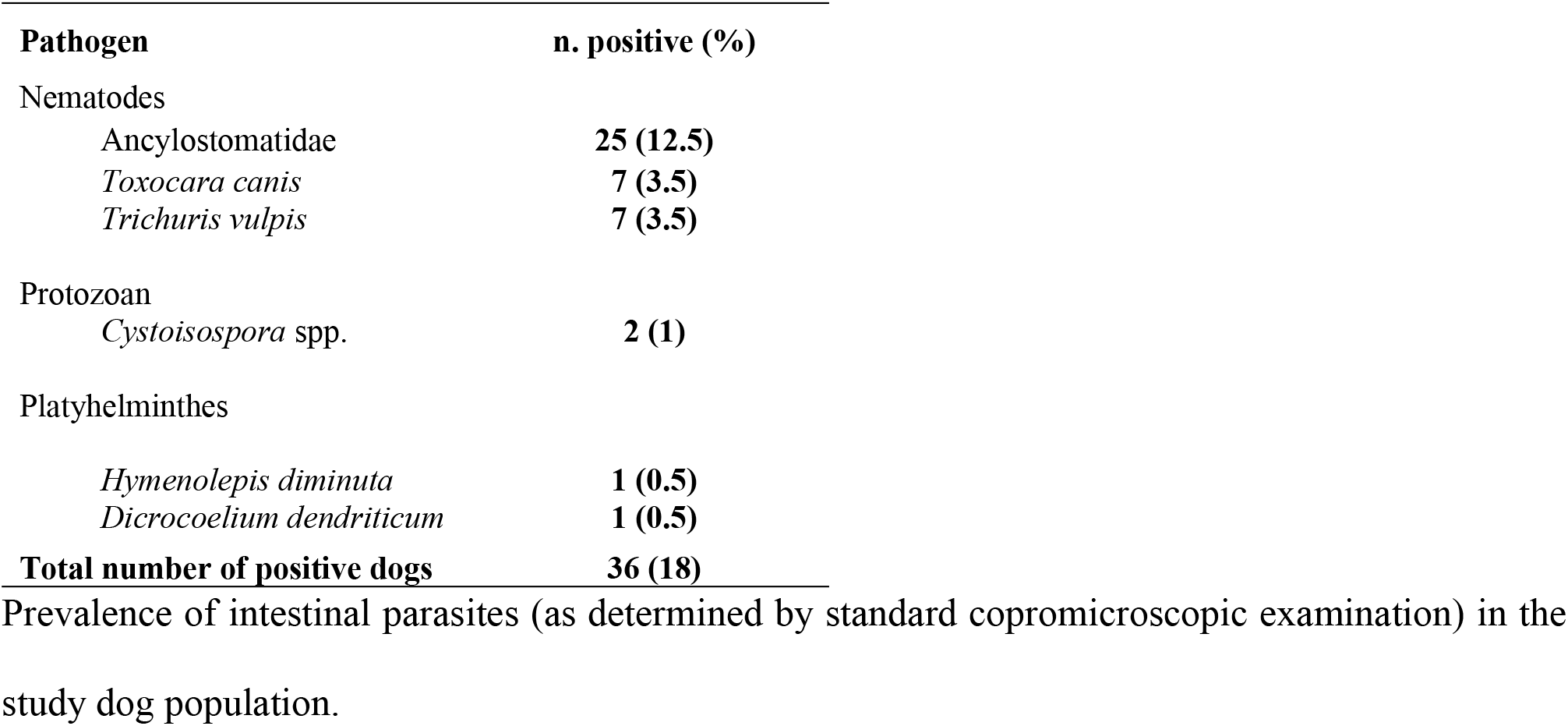

**Table 4.**

| Variable | Positive for<br>intestinal parasites<br>Odds ratio<br>(95% CI)<br>*GLM P-value* | Positive for<br>VBDs<br>Odds ratio<br>(95% CI)<br>*GLM P-value* | Positive for<br>filariae<br>Odds ratio<br>(95% CI)<br>*GLM P-value* | Positive for<br>ectoparasites<br>Odds ratio<br>(95% CI)<br>*GLM P-value* | Zoonotic<br>infections<br>Odds ratio<br>(95% CI)<br>*GLM P-value* |
| --- | --- | --- | --- | --- | --- |
| *Island* |  |  |  |  |  |
| Santorini vs Tinos | 5.49<br>(1.60-18.83)<br>**0.007\*\*** | 0.30<br>(0.13-0.67)<br>**0.005\*\*** | Na | 0.58<br>(0.24-1.37)<br>0.211 | 0.66<br>(0.28-1.59)<br>0.357 |
| Santorini vs Ios | 11.43<br>(1.31-99.56)<br>**0.027\*** | 0.76<br>(0.31-1.85)<br>0.548 | Na | 4.51<br>(1.52-13.36)<br>**0.007\*\*** | 1.73<br>(0.70-4.30)<br>0.233 |
| Santorini vs Skiathos | 23.86<br>(5.51-103.33)<br>**<0.001\*\*\*** | 0.24<br>(0.10-0.60)<br>**0.002\*\*** | Na | 19.28<br>(5.51-67.41)<br>**<0.001\*\*\*** | 0.77<br>(0.29-2.07)<br>0.613 |
| *Age* |  |  |  |  |  |
| Up to 1 yr vs 1 to 7<br>yrs | 0.25<br>(0.06-0.97)<br>**0.046\*** | 0.36<br>(0.14-0.96)<br>**0.040\*** | Na | 0.67<br>(0.24-1.89)<br>0.458 | 0.26<br>(0.10-0.69)<br>**0.007\*\*** |
| Up to 1 yr vs more<br>than 7 yrs | 0.26<br>(0.05-1.35) 0.108 | 0.54<br>(0.18-1.64)<br>0.277 | Na | 0.59<br>(0.17-1.97)<br>0.392 | 0.51<br>(0.17-1.54)<br>0.229 |
| *Sex* |  |  |  |  |  |
| Male vs Female | 1.36<br>(0.54-3.45)<br>0.518 | 2.33<br>(0.24-4.37)<br>**0.008\*\*** | 0.61<br>(0.07-5.45)<br>0.656 | 0.72<br>(0.36-1.45)<br>0.356 | 2.13<br>(1.11-4.10)<br>**0.023\*** |
| *Lifestyle* |  |  |  |  |  |
| Stray vs Owned | 11.37<br>(3.73-34.72)<br>**<0.001\*\*\*** | 1.15<br>(0.55-2.40)<br>0.707 | 0.29<br>(0.03-2.98)<br>0.300 | 3.17<br>(1.37-7.34)<br>**0.007\*\*** | 3.00<br>(1.37-6.60)<br>**0.006\*\*** |
| *Travel* |  |  |  |  |  |
| Yes vs No | 0.49<br>(0.05-4.48)<br>0.526 | 1.34<br>(0.58-3.11)<br>0.498 | Na | 0.33<br>(0.11-0.97)<br>**0.043\*** | 1.10<br>(0.48-2.53)<br>0.819 |

Statistical analysis evaluating various factors (i.e. island where the dogs lived, age, sex, lifestyle and traveling history) in relation to the different infections detected in the study animals. \*Statistical significant result at the 0.05 level; \*\*Statistical significant result at the 0.01 level; \*\*\* Statistical significant result at the 0.001 level.

##### Ectoparasites

Fleas were recorded in 26.5% of the dogs (53/200), specifically in 24.2% (16/66), 58% (29/50), 9.3% (4/43) and 9.8% (4/41) dogs from Santorini, Tinos, Ios and Skiathos respectively. All collected specimens were identified as *Ctenocephalides felis felis*. Ticks were present in 25% of the dogs (50/200), namely in 40.9% (27/66), 38% (19/50) and 9.3% (4/43) from Santorini, Tinos and Ios respectively. All ticks were *Rhipicephalus sanguineus sensu latu*. Mites were not found in any of the investigated dogs.

Mixed infections by ticks and transmitted diseases were recorded in 28 (14%) dogs while no dogs showed mixed infection by fleas and transmitted pathogens.

The occurrence of fleas and ticks was statistically associated with study site, lifestyle and travelling. Specifically, a higher risk of ectoparasitosis was recorded in Santorini against Ios (odds ratio=4.51) and Skiathos (odds ratio=19.28), while the same risk was found in Tinos. Stray dogs increased the odds of ectoparasitosis (odds ratio=3.17), while travelling decreased the odds (odds ratio=0.33) (Table 4).

#### Blood examination (Table 5)

##### Rickettsia conorii

Overall, 93 dogs (46.5%) seroreacted to *R. conorii*. Twenty-two (11%) were also PCR positive and, at the blood smear examination, the samples of seven (3.5%) of them was positive for corpuscoles suggestive *Rickettsia* spp. in the monocytes. Sequences from the 22 PCR products showed 99% identity with *R. conorii* GenBank Accession number U59730.1. All dogs negative by serological analyses were also negative for *Rickettsia* spp. upon PCRs. Of the 93 *R. conorii-* seropositive dogs, 19 had a tick infection and 6 scored also PCR-positive for *R. conorii*. The prevalence of *R. conorii* in animals was found statistically related with the presence of lymphadenopathy (Fisher’s exact test, *p*=0.0177) whilst no any relation was detected with signs like weight loss, pale mucous membrane, gastrointestinal disorders, conjunctivitis or dermatological manifestations (Fisher’s exact test, *p*≥0.05).

**Table 5.**

| Pathogen | PCR | Serological<br>detection | Smear or other direct<br>parasitological examination |  |
| --- | --- | --- | --- | --- |
|  | n. positive (%) | n. positive (%) | n. positive (%) | n. positive (%) by at<br>least one of the test |
| Anaplasmataceae |  |  |  |  |
| *Ehrlichia canis* | 3 (1.5) | 19 (9.5) | 0 | 19 (9.5) |
| *Anaplasma* spp. | 1 (0.5) | 2 (1) | 1 | 2 (1) |
|  | *A. ph.* | *A. ph/A.pl* | *Anaplasma* spp. | *A. ph/A.pl* |
| *Leishmania infantum* | 13 (6.5) | 13 (6.5) | 0 | 13 (6.5) |
|  | *L. i.* | *L. i.* |  | *L. i.* |
| *Rickettsia conorii* | 22 (11) | 93 (46.5) | 7 | 93 (46.5) |
|  | *R. c.* | *R. c.* | *Rickettsia* spp. | *R. c.* |
| *Babesia canis* | 1 (0.5) | 1 (0.5) | 0 | 1 (0.5) |
|  | *B. c.* | *B. c.* |  | *B. c.* |
| *Dirofilaria* spp. | ND | 1 (0.5) | 1 (0.5) | 1 (0.5) |
|  |  | *D. i.*<br>ND | *D. i.*<br>3 (1.5) | *D. i.*<br>3 (1.5) |
|  |  |  | *D. r.* | *D. r.* |
| *Borrelia burgdorferi* | ND | 0/200 (0) | ND | 0/200 (0) |

Observed prevalence of vectorn borne pathogens (as determined by PCR, serology and blood microscopy) in the study dog population. *A. ph.*: *Anaplasma phagocytophilum*; *A. pl.*: *Anaplasma platys*; *L. i.*: *Leishmania infantum*; *R. c.*: *Rickettsia conorii*; *B. c.*: *Babesia canis*; *D. i.*: *Dirofilaria immitis*; *D. r.*: *Dirofilaria repens*.

##### Anaplasmataceae

Nineteen dogs (9.5%) were seropositive for *E. canis*, three of them being also positive at the PCR (100% homology with Genbank Accession number LC018188.1), while the others were PCR-negative. Two dogs (1%) seroreacted for *A. phagocytophilum/A. platys*. One of them showed corpuscoles suggestive of *Anaplasma* spp. in the monocytes at the Giemsa staining and scored PCR-positive for *A. phagocytophilum* (homology of 100% with Genbank Accession number KY114936.1), while the other was microscopically and PCR-negative.

Ticks were present in 15/19 dogs seropositive for *E. canis* at the SNAP 4Dx Plus and 3 of them were among those that scored positive at the PCR.

##### *Babesia* spp

One dog (0.5%) was serologically and PCR positive for *B. canis* showing homology of 99% with GenBank Accession number: KJ696714.1, while all other dogs were negative.

##### Leishmania infantum

A total of 13 (6.5%) samples seroreacted for *L. infantum* and all of them were also PCR-sequencing positive, with 99-100% identity with *L. infantum* GenBank Accession number HM807524.1. All seronegative dogs were also negative upon PCR.

##### *Dirofilaria* spp

One (0.5%) of the examined dogs was positive for *D. immitis*, at both Knott’s test and serology. Moreover, in three dogs (1.5%) microfilariae of *Dirofilaria repens* were found in the Knott’s test.

##### Risk factor for exposure to parasites and VBDs

The percentage of dogs positive for at least one intestinal parasites and/or VBDs was similar between travelling (i.e. 58.3%) and non-travelling dogs (66.5%) while the highest number of infected animals was found in sheltered animals (83.1%).

The exposure to at least one VBD was statistically associated to the study site, the sex and the age. In particular, animals residing in Skiathos were 4.17 (i.e. 1/0.24) and 3.17 (i.e. 0.76/0.24) times more likely to be infected than those in Santorini and Ios, respectively, while no difference was found compared to dogs from Tinos. Furthermore, VBD prevalence was associated with sex and age, as males increased the odds (odds ratio=2.33) and dogs ageing up to 1 year decreased the odds (odds ratio=0.36) against those ageing 1-7 years.

The diagnosis of VBDs was associated with the presence of various skin lesions (GLM, *p* <0.05) in infected animals. In fact, animals with skin lesion were 3.2 times more likely to be seropositive compared to those with no dermatological manifestations. Similarly, weight loss was associated with 5.43-fold increased odds (Table 6). Nonetheless, positivity to a VBD was not related to the presence of ectoparasites (Fisher’s exact test, *p*= 0.6623), as clinical signs (i.e. lymphadenopathy, pale mucous membrane or conjunctivitis) were not statistically significant factors (GLM, *p*>0.05).

**Table 6.**

|  |  |  |
| --- | --- | --- |
|  | Positive for VBD | Positive for *L. infantum* |

| Variable | Odds ratio<br>(95% CI) | GLM<br>P-Value | Odds ratio<br>(95% CI) | GLM<br>P-Value |
| --- | --- | --- | --- | --- |
| *Lymphadenopathy* | 0.66 | 0.335 | 0.82 | 0.815 |
| Yes vs No | (0.28-1.54) |  | (0.16-4.30) |  |
| *Dermatological manifestations* | 3.20 | **0.027\*** | 2.62 | 0.205 |
| Yes vs No | (1.14-8.97) |  | (0.59-11.59) |  |
| Weight loss | 5.43 | **0.033\*** | 2.87 | 0.210 |
| Yes vs No | (1.15-25.65) |  | (0.55-14.90) |  |
| *Pale mucous membrane* | 0.48 | 0.242 | 0.62 | 0.679 |
| Yes vs No | (0.14-1.65) |  | (0.06-6.06) |  |
| *Conjunctivitis/Ocular manifestations* | 4.86 | 0.154 | Na | 0.992 |
| Yes vs No | (0.55-42.54) |  |  |  |

Statistical analysis to evaluate the presence of clinical signs in relation to the different infections detected in the examined dogs.

#### Geographic distribution of parasites

Santorini showed the highest prevalence of infected dogs (65.2%) and the highest value of infection by Ancylostomatidae (27.3%) (Fisher’s exact test, *p*<0.001), while Skiathos had the highest rate of dogs that seroreacted to *R. conorii* (73.2%) (Fisher’s exact test,*p*<0.001) and of dogs infected by *D. immitis* (2.4%) and *D. repens* (7.3%) (Fisher’s exact test, *p*=0.008). Skiathos showed the highest percentage of positivity to *L. infantum* (17.1%).

#### Zoonotic risk

The risk to carry zoonotic pathogens (i.e. *R. conorii, Anaplasma* spp., *L. infantum, Dirofilaria* spp., Ancylostomatidae and *T. canis*) was associated to animal age, as dogs ageing 1-7 years increased the odds of infections by 3.85-fold (i.e. 1/0.26) in comparison to younger animals. Furthermore, the sex and lifestyle were also found statistically related with zoonotic infections with male and stray dogs being 2.13 and 3 times more likely affected than female and owned dogs, respectively.

### Discussion

In the present study 65% of the dogs showed exposure to or were carrying pathogens, and 64% harboured a pathogen with a zoonotic potential. Among them, canine geohelminths have an important health impact for both animals and humans. Thus, dogs here found infected by *T. canis* or hookworms represent a potential health risk especially because free roaming animals are a source of environmental contamination [25]. Human infection by *T. canis* may lead either to subclinical infections or to different *larva migrans* syndromes i.e. visceral, ocular and neural, that may have serious clinical manifestations [26, 27]. *Ancylostoma caninum* larvae may penetrate the human skin and cause follicular, papular/pustule and ephemeral lesions, muscular damages and, seldom, eosinophilic enteritis. The risk of human infection is enhanced by walking barefoot or lying on grass and soil in contaminated areas [27].

Eggs of the tapeworm *H. diminuta* were found in one dog. Althoug the main hosts of this parasite are rodents [28, 29], in rare cases dogs and other mammals, including humans, may also be infected. Similarly, eggs of *D. dentriticum* a trematode commonly found in the liver of ruminants, were found in one dog. While in some cases this parasite can infect other mammals, including dogs and humans [30, 31], it is plausible that pseudoparasitism is the cause of this finding [32].

The high rate of exposure of the here studied dogs to several pathogens transmitted by arthropods, is of importance because some VBDs are shared between companion animals and people. Moreover, to the best of the authors’ knowledge, this is the first report of seroprevalence and molecular detection of *Babesia* spp., *E. canis, R. conorii* and *Anaplasma* spp. in dogs from the herein study areas.

The high seroprevalence for *R. conorii*, i.e. the main aetiological agent of the Mediterranean spotted fever in Europe [33], suggests a frequent exposure and/or persistent low-grade infections. This is not surprising if one considers the geographical location of the study sites, i.e. regions in Southern Europe with favorable environments for ticks and local circulation of transmitted pathogens. It is worth noting that the DNA of this bacterium was found in 11% of examined dogs, thus further corroborating recent findings that have indicate dogs as important reservoirs of this pathogen and source of infection for ticks [34–36]. The role of dogs as reservoirs is further supported by the absence of clinical signs or by the presence of mild aspecific alterations, as in the present study (e.g. lymphadenopathy). Under an epizootiological standpoint it is remarkable that all ticks collected from positive animals were *R. sanguineus*, i.e. the main vector of *R. conorii* [33]. It is also interesting that a past survey carried out in the island of Crete showed a human seroprevalence rate of 7.6% for *R. conorii* [37]. Although no ticks from Crete were PCR-positive for *R. conorii* [37], the bacterium was isolated from *R. sanguineus* and humans in other areas of Greece with high seroprevalence in people [38, 39]. More importantly, several cases of human Mediterranean spotted fever have been reported in Greece [40, 41].

*Ehrlichia canis* causes the canine monocytic ehrlichiosis, a severe and potentially life-threatening illness in dogs transmitted in Europe by *R. sanguineus* [42–44]. Altough an *E. canis-like* organism has been described in humans in Venezuela [45], to date this VBD is not considered zoonotic. The infection rate by *E. canis*, much lower than that recorded for *R. conorii*, could appear unexpected, but it is similar with that recorded in a recent study from southern Italy [46]. It should be noted that different lineages of *R. sanguineus* have a low capacity to transmit *E. canis* [47], thus explaining why in some cases the prevalence of *E. canis* infection could result lower than expected [48, 49].

It could be hypothesized that some *R. sanguineous* lineages may have opposite ability to transmit either *R. conorii* or *Ehrlichia/Anaplasma*. Further studies to investigate this issue are thus warranted. *Anaplasma phagocytophilum* was detected in one dog. It is an emerging vector-borne pathogen transmitted by *Ixodes ricinus* ticks [50]. This tick species was not found on the dogs examined in the present study. However, it is widespread in continental Greece [49, 51, 52] and further investigations to evaluate its presence in insular Greece could be thus useful. The absence or limited occurrence of *A. platys*, the agent of infectious cyclic thrombocytopenia, is more surprising as this bacterium is probably transmitted by *R. sanguineus* ticks and common in the Mediterranean regions [43, 53]. Again, this epidemiological feature could be due to the presence of different lineages of the brown dog tick.

Although the most common canine haemoprotozoan in Mediterranean countries is *Babesia vogeli, B. canis* is also enzootic in Europe [3], including in countries neighbouring to Greece and the Balkan Peninsula [54, 55]. The presence of *B. canis* in insular regions of Greece is an unexpected result because its primary vector, *Dermacentor reticulatus*, generally lives in cool and wet climates. As *D. reticulatus* can also be found in warm and temperate areas [56], its sporadic presence in Greece, especially in continental regions, needs to be taken into account. The owner of the dog declared that the dog had travelled in continental Greece but further details were not provided. Thus, the origin of this result could be likely due to an infection acquired in other areas of the Country, although further information was not available.

The presence of several *L. infantum-positive* dogs in all study sites is explained by its wide distribution in Southern Europe [57], including Greece where mean seropositivity is around 20% [5, 6]. This high infection pressure is shown also by the recent records of infection in cats from Crete and Athens [9] and by the more than 300 autochthonous human cases reported in 2005-2010 according the Hellenic Center for Disease Control and Prevention [6]. The zoonotic potential of *L. infantum* in humans is of great importance, as the disease may be severe and presents with visceral, cutaneous and mucocutaneous signs [58].

Dogs living in Skiathos proved to be at risk of dirofilariosis. Though *Dirofilaria* spp. have been here diagnosed with a low prevalence, their ability in causing disease in humans should be taken into account. The number of human cases of *D. repens* infection in Europe is currently a public health concern and the lack of awareness with diagnosis and control in microfilariaemic dogs could lead to lack of vigilance and underestimation for this parasite [59]. *Dirofilaria* infections were here found only in Skiathos (where endemic infections by *D. immitis* is absent) in a single dog that was imported from an area of Central Greece where the prevalence of canine dirofilariosis is about 7% [60]. This confirms that undiagnosed and untreated microfilariaemic dogs are an important source of infection for mosquitoes and may also introduce these pathogens in other regions.

The association of the study sites with given infections may be due to the different island characteristics. Skiathos, where dogs resulted more likely infected with VBDs and ectoparasites than in other sites, is a part of the Vories Sporades, an island formation with a rich vegetation that fosters the maintenance of ectoparasites and vectors more efficiently. Indeed, the dry environment of Santorini, Ios and Tinos, belonging to different island formations (i.e. Cyclades) is characterized by poor vegetation.

The higher occurrence of intestinal parasites in Santorini rather in the other islands is difficult to explain. These results could have originated by chance but it could be hypothesized that differences in regular epizootiological vigilance and appropriate veterinary care in terms of prevention, diagnosis and treatment have a certain role, although further investigations are warranted to clarify these issues. The higher occurrence of intestinal nematodes in dogs ageing 1-7 years should be interpreted with caution given the lack of precise information provided in some cases by the owners about anthelmintic treatments. Owners could be less willing to engage in preventative or therapeutic practices for dogs ageing more than 1 year because erroneously considered at less risk of intestinal parasitoses. These data further indicate that a high level of vigilance comprising a routine fecal examinations and appropriate anthelmintic treatments are still indicated in adult dogs and should be encouraged among owners.

In conclusion, it is here shown that regular investigations should be encouraged in areas where data on parasite occurrence is still limited, and that prompt diagnosis and infection control are a prority. This is especially relevant for Mediterranean countries where epizootiological and biological conditions favour the occurrence and circulation of several parasitoses and VBDs in animal populations due to good and animal trade, dog adoptions and holiday trips of people [36, 43, 61–63]. As visitors and tourists often bring their pets during holidays, introduction of parasites from endemic to free areas can occur. At the same time there is the risk that these dogs acquire pathogens in a new, enzootic environment and introduce them in free areas when they go home. In this view, timely use of dewormers (e.g. macrocyclic lactones) secures treatment of zoonotic geohelminthoses and at the same time provide efficacious prevention for spreading (e.g. *D. immitis*) or largely distributed (e.g. roundworms) parasites. Also, the appropriate use of acaricides, insecticides and repellents are reliable ways to improve dog health and to control environmental infection by vectors, preventing the spread of potentially deadly diseases.

## Acknowledgements

The authors thank Carmine Merola and Raffaella Iorio for the participation in this study. Special acknowledgments are due to the veterinarians in the Islands of the study: Maria Danezi, Kalliopi Tsagari and Manina Vlastaridou as well as to the Greek Action of Volunteer Veterinarians (GAVV) for their valuable help and excellent collaboration.

